# Sprod for De-noising Spatial Transcriptomics Data Based on Position and Image Information

**DOI:** 10.1101/2021.11.03.467103

**Authors:** Yunguan Wang, Bing Song, Shidan Wang, Mingyi Chen, Yang Xie, Guanghua Xiao, Li Wang, Tao Wang

## Abstract

Spatial transcriptomics (ST) technologies provide gene expression close to or even superior to single-cell resolution while retaining the physical locations of sequencing and often also providing matched pathology images. However, the expression data captured by ST technologies suffer from high noise levels, as a result of the shallow coverage in each sequencing unit. The extra experimental steps for preserving the spatial locations of sequencing could result in even more severe noises, compared to regular single-cell RNA-sequencing (scRNA-seq). Fortunately, such noises could be largely removed by leveraging information from the physical locations of sequencing, and the tissue and cellular organization reflected by corresponding pathology images. In this work, we demonstrated the extensive levels of noise in ST data. We developed a mathematical model, named Sprod, to impute accurate ST gene expression based on latent space and graph learning of matched location and imaging data. We comprehensively validated Sprod and demonstrated its advantages over prior methods for removing drop-outs in scRNA-seq data. We further showed that, after adequate imputation by Sprod, differential expression analyses, pseudotime analyses, and cell-to-cell interaction inferences yield significantly more informative results. Overall, we envision denoising by Sprod to become a key first step to empower ST technologies for biomedical discoveries and innovations.

## INTRODUCTION

Spatially resolved transcriptomics (ST) technologies have recently bloomed, such as 10X Visium, Slide-Seq^1,2^, HDST^3^, Seq-scope^4^, XYZeq^5^, *etc*. STs provide gene expression information for the whole transcriptome at near- or sub-single cell resolution, in tandem with matching spatial information and, for many techniques, also the matched pathology images stained by Hematoxylin and eosin (H&E) or immunofluorescence (IF). The units of sequencing are called spots or beads by different techniques but are essentially a small cluster of cells (up to a few dozens), or even parts of a single cell for some newer techniques with higher spatial resolution, such as HDST. These powerful techniques have enabled researchers to localize cell types within the tissues, to characterize spatial expression patterns, to define the local ecosystems of cells, and to resolve the spatiotemporal order of cellular development.

However, such technologies suffer from severe noises in the gene expression measurements. The noises come from random variations due to the shallow nature of sequencing for each spot/bead, similar to regular single-cell RNA-sequencing (scRNA-seq). But they are also further complicated by the extra and delicate experimental steps needed to preserve the spatial locations of sequencing. Thus, pre-processing steps of ST data are necessary before any downstream analyses, in order to remove such noises. The methodologies developed for addressing scRNA-Seq drop-outs (loss of expression)^6,7^ are likely insufficient for this purpose, as the expressional noises in STs are more than just drop-outs. Moreover, such methods only rely on the expression data themselves to correct for drop-outs, and such “bootstrapping” methodologies are limited in the extent to which the drop-outs can be reliably corrected. In other words, they ignored the spatial and imaging features of the spots/beads provided by STs, which can potentially guide and improve the correction of noises in STs with useful external information.

In this work, we demonstrated the existence of extensive noises in the ST data. We developed Sprod, short for **S**patial **Pro**filing **D**e-noising, to impute accurate gene expression, by leveraging the location information of each measurement, and the corresponding imaging data, which are available for most of ST techniques. By testing on a variety of ST datasets, we systematically validated Sprod’s accuracy and robustness. We also showed its superiority to existing drop-out removal methods of scRNA-seq data. With Sprod-corrected ST data, we discovered 3-4 times more RNA transcripts that were transported from the soma of the neurons into the dendrites in the mouse hippocampus, compared to the un-corrected ST data. With Sprod, we also better delineated the different transcriptomic features of the tumor cells near the boundaries of the tumor-stroma/immune regions compared with those in the centers of the tumor regions, *via* pseudotime analyses. These differences were shown to be driven by tumor cell-stroma/immune cell interactions at their interfaces. Overall, careful handling of technical noises in ST data is a critical first step to the unbiased discovery of new biological knowledge.

## RESULTS

### Extensive Noises Exist in ST Data

We demonstrated the existence of extensive noises in ST data. First, we investigated a 10X Visium ovarian cancer dataset, with matched immunofluorescence (IF) images of CD45/LCA (Leukocyte Common Antigen), keratin, and DNA. **Fig. 1a** presented the CD45 protein expression obtained by IF staining, and the RNA expression level of *PTPRC* (the gene encoding CD45), obtained from both the target panel sequencing and whole transcriptome sequencing by Visium. Strikingly, there is a very poor correlation between CD45 protein IF and *PTPRC* RNA expression (by either target panel or whole transcriptome). When the spots were subset by keeping only those with higher overall sequencing depth, which are potentially of higher quality, the correlation improved by a large extent (**Sup. Fig. 1**). One of the many possible sources of inaccuracies results from drop-outs, as in regular scRNA-seq^6–8^. In **Fig. 1b**, it is clear that severe drop-outs exist in the expression of *PTPRC* (excessive zeros at X=0). We created a gene signature of *PTPRC* RNA expression by including highly correlated genes of *PTPRC*, which essentially removed drop-outs by *ad hoc* averaging. We found that this *PTPRC* RNA signature’s correlation with CD45 protein IF intensity is further improved (**Sup. Fig. 1**), compared with the single gene of *PTPRC*.

**Fig. 1.**
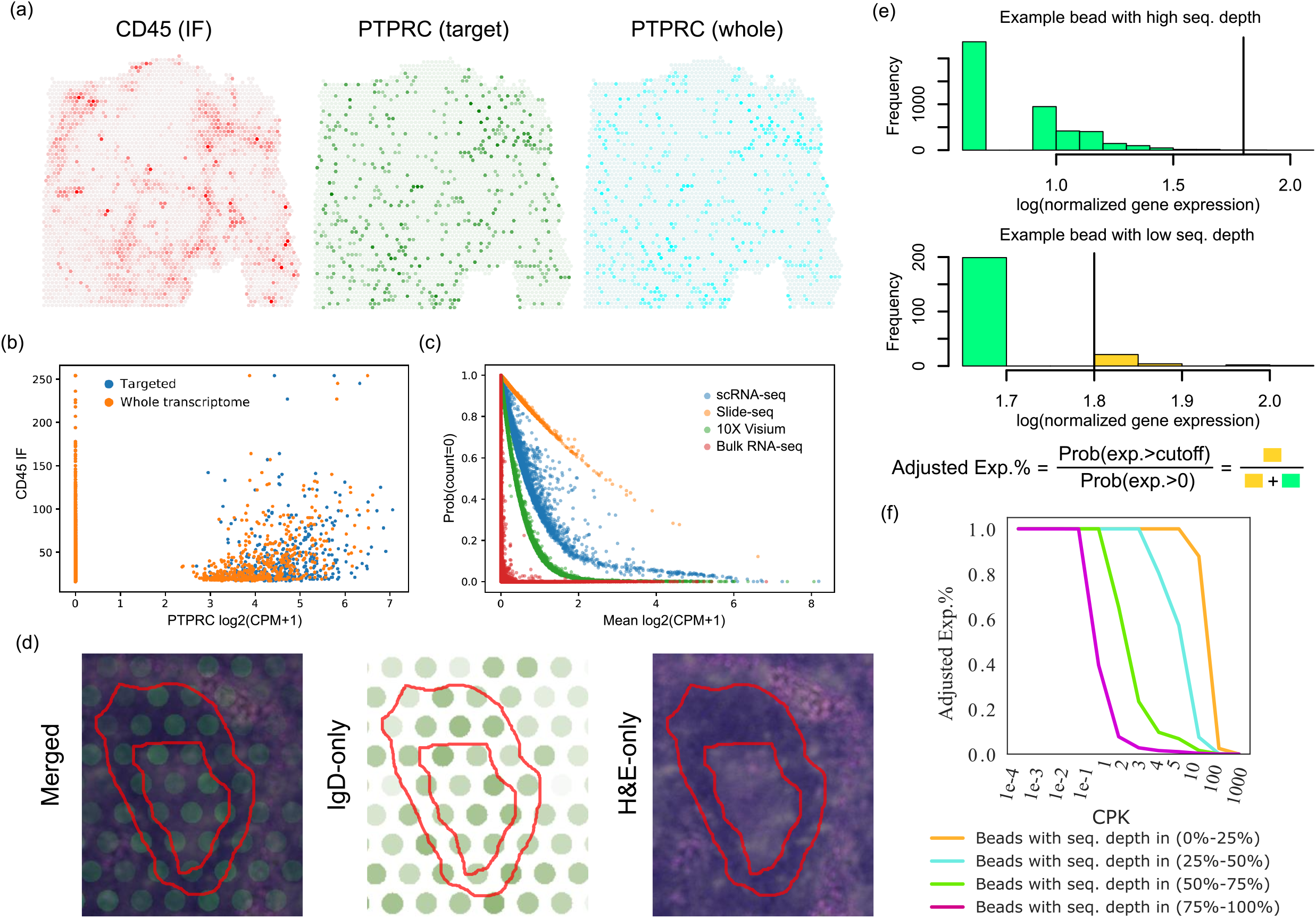
Extensive noise in spatial transcriptomics data. (a) The spatial expression levels of CD45 (IF), *PTPRC* (targeted panel sequencing) and *PTPRC* (whole transcriptomic sequencing) of the 10X Visium Ovarian Cancer dataset. (b) Severe drop-outs in the *PTPRC* gene expression for both the targeted sequencing and whole transcriptomic sequencing. X-axis shows the *PTPRC* RNA expression level (unit=log2(CPM+1)). (c) Severe drop-outs in scRNA-seq, Slide-Seq, Visium and bulk RNA-Seq. The X-axis is the average RNA expression levels of each gene profiled by each technique/dataset, and the Y axis shows the percentages of counts of exactly 0 for each gene. (d) One example mantle zone structure with poor agreement with IgD expression from the 10X Visium human lymph node dataset. The red lines mark the borders of the mantle zone. (e) A cartoon describing the schema of the calculation of Adjusted Exp.%. (f) The Adjusted Exp.% of beads in the four bins of different sequencing qualities, in the mouse Slide-Seq dataset, and with the cutoffs to define the Adjusted Exp.% shown on the X axis.

Indeed, we observed that drop-outs are a severe issue in both Visium and Slide-seq datasets. In **Fig. 1c**, we showed the percentages of zero counts of all captured genes, as a function of the average expression of each individual gene across all beads/spots, for expression measurements from Visium, Slide-seq, regular scRNA-seq, and regular bulk RNA-seq (from TCGA breast cancer cohort). In all techniques, the percentages of 0 counts increased with lower average gene expression levels, as expected. Visium and Slide-seq both have drop-out levels comparable to regular scRNA-seq. Visium’s drop-outs rates are lower than regular scRNA-seq, as each spot of Visium sequencing usually contains a group of cells. Slide-seq has a much higher resolution (very close to single-cell resolution), and a much higher rate of drop-outs than regular scRNA-seq, possibly due to its lower per-cell sequencing depth.

However, the noises from ST data can also be an inflation of gene expression, rather than only a loss of expression (extreme cases are drop-outs). We examined another 10X Visium dataset of human benign reactive lymph nodes. In the normal lymph nodes, follicles and perifollicular “ring-like” structures exist (the darker rings in the H&E stained tissues images of **Fig. 1d** and **Sup. Fig. 2**), which represent reactive germinal centers (GCs) surrounded by well-defined mantle zones^9–11^. However, examination of the expression of IgD, a marker of mantle zone, in this Visium dataset shows only a weak correlation with the ring-like mantle zone structures on the H&E stained tissue sides. In one example mantle zone (**Fig. 1d**), the green color, which stains IgD expression, only weakly correlates with the purple staining, which indicates the mantle zone. More importantly, drop-outs can not completely explain the observed weak correlation, as it appears that IgD expressions are strong in some places (such as the GC in the center of this mantle zone) where they should not be.

The Slide-Seq data was also investigated. Slide-Seq data do not have the matched imaging data for us to cross-reference as a gold standard. Therefore, we investigated another aspect of the Slide-Seq data. We noticed that Slide-Seq has a much shallower sequencing depth compared with Visium and regular scRNA-Seq. This could result in severe noises in the gene expression measurement, where the measured expression could be either artificially higher or lower than the truth. Here we focused on the problem of inflated high expression (false positive), as the problem of drop-outs (false negative) has already been investigated in our previous example. We hypothesized that the beads with lower sequencing quality are more likely to demonstrate overly inflated expression. As we showed in **Sup. Fig. 1** that the total sequencing depth of each bead can be used as a surrogate of bead sequencing quality, we divided all beads into 4 bins, based on the total sequencing depth. Then for each gene, we investigated the probability that the beads would show a higher expression than a given cutoff in each of the bins, among the beads that have the non-zero expression of this gene (**Fig. 1e**). Since the bead sequencing quality is a technical issue, we expected the beads in different bins to follow similar distributions of gene expression. However, as we showed in **Fig. 1f**, there is a dramatic difference between beads in different quality bins in terms of their gene expression distribution. The bins with lower quality are more likely, for all cutoffs employed (X-axis), to yield expression counts that are larger than the cutoffs (Y-axis). This unexpected distributional difference among bins of different technical qualities could confer a high level of bias in gene expression analyses.

### Sprod for De-noising ST Data Based on Latent Graph Learning

To remove the significant level of noises in the ST expression data, we developed Sprod (**Fig. 2a**), which corrects the noises in expression data guided by the location and imaging information. Sprod operates in two stages. In the first stage, Sprod leverages the spatial locations of the beads/spots to determine the neighborhoods of the spots/beads to borrow information. However, it is imperative to consider the cell type heterogeneity of the spots/beads in this neighborhood. Therefore, borrowing information is only restricted to the cells of the same type. For the Visium platform, ST data are provided with the matched pathology images, from which textures and channel intensities can be derived to inform the cell type heterogeneity. On other hand, for the Slide-Seq platform, no matched images are provided. We leverage the fact that the overall transcriptomic profiles of the beads should be sufficient for a simple cell type clustering task, which essentially averages out individual genes’ noises in an *ad hoc* manner. Therefore, we use the overall gene expression profiles of the beads to perform unsupervised clustering to detect different types of cells. Then Sprod creates a “pseudo-image”, whose “image channels” are essentially the detected cell clusters/types and their assignment probabilities. In the second stage of the model, de-noised gene expression is generated by capturing the local information on the manifold of the learned similarity graph, namely by borrowing information from expression data across beads/spots through the graph edges. The technical details of Sprod are specified in the method section and **Sup. File 1**.

**Fig. 2.** Sprod for de-noising of spatial transcriptomics data. (a) A cartoon describing the schema of the Sprod model, from data preparation, graph building, to expressional de-noising. (b) The simulated dataset. The figure shows the location of the simulated spots, and the cell types they are assigned to. The light gray bars show the graph built by Sprod. A bar connects two dots as long as the graph connects two spots, regardless of edge weights. However, the dots have been subsampled to avoid over-cluttering of the presentation. (c) The denoising% of all parameter combinations tested. The X axis shows all parameter combinations, ordered from lower to higher denoising%. (d) Visualizing the denoising% with respect to specific choices of parameters. K=10. The X and Y axes show all the choices tried for each of R and Lambda. The Z axis shows the denoising%, which is the average of the denoising% over all the other parameters’ choices. Noise level=0.5. (e) An example of the two diagnostic plots generated by Sprod from the 10X breast cancer Visium dataset. Left, the spots and edges of the detected graph on the X-Y coordinates. Right, the spots and edges of the detected graph in the Umap space of the image features. The coloring denotes confidence of the edges, with blue referring to high confidence and gray referring to low confidence.

We performed simulation analysis to evaluate and showcase the performance of Sprod. We simulated a dataset of 5,000 spots (**Sup. File 2**). The spots were divided into three cell types, A, B, and C. The details of the simulation were specified in the method section. We applied Sprod to this simulation dataset. We first checked the graph of spatial/image similarity constructed by Sprod and found that the graph generally correctly connected spots of the same cell types that are also in spatial proximity (one example shown in **Fig. 2b**). We evaluated the denoising performance using the sum of absolute error (SAE) between the denoised/raw expression and the clean expression (with no simulated noise). The metric for our evaluation is defined as denoising%=1-SAE(denoised)/SAE(raw). We tried the combinations of all tuning parameters on this dataset (all tested parameters described in the method section). **Fig. 2c** showed that Sprod can decently remove the noises in the ST data across a wide range of parameter combinations. Our exploration of these parameter combinations shows that three tuning parameters, R (the radius to define the neighborhood of the spots), K (the dimension of the latent space), and Lambda (a scaling parameter to control clustering sparsity), have a critical contribution to the denoising%, while the other tuning parameters (such as the tSNE-type perplexity for building the similarity graph) have minimum influences. Overall, K=10 works the best. In **Fig. 2d**, we showed a surface plot of the performances of Sprod (denoising%) with respect to the choices of R and Lambda, given K=10, which stably exceeds 80%. We arrived at a set of optimal tuning parameters for this simulation dataset (**Sup. Table 1**), which serves as a reference for the application of Sprod on the real datasets below.

In practice, the simulation data are still different from the real datasets, and real datasets are also different from each other, inevitably calling for tuning of the R, K, and Lambda parameters. We recommend the users to start from the optimal parameter set in our simulation dataset above and use two sets of diagnostic plots that we provided in the Sprod software (**Fig. 2e**, the 10X breast cancer dataset shown as an example) to choose the optimal parameter set. The first diagnostic plot displays the spots/beads and the edges of the similarity graph on the physical X-Y coordinates, which can inform whether the graph has correctly captured the pattern of tissue organization. With a good parameter set, the pattern of the similarity graph edges should be reflective of the pattern of the pathological images. The other two diagnostic plots display the spots/beads and the similarity graph in the imaging feature space (dimension reduction performed through t-SNE and uMap; the uMap result in **Fig 2e**). With a good parameter set, the similarity graph edges will connect the spots/beads that are close to each other in the imaging feature t-SNE/uMap plots. While **Fig. 2e** shows example diagnostic plots of “good” parameters, we also showed in **Sup. File 2** some example diagnostic plots of “bad” parameters, to help users understand how to choose tuning parameters for Sprod.

### Validation of Sprod Shows Its capability to Remove the Noises in ST Data

Sprod was first applied on a 10X Visium ovarian cancer dataset. For the matched images, we included the Keratin and DAPI IF channels and left out the CD45 IF channel for independent validation. **Fig. 3a** shows the denoised *PTPRC* expression on the slide. The “discrete” appearances of the original *PTPRC* expression (**Fig. 1a**) disappeared and the adjusted *PTPRC* expression demonstrates a more continuously changing pattern, and more importantly, is consistent with the pattern of CD45 IF intensities (**Fig. 1a**). We showed a scatterplot of the CD45 IF intensities and the denoised *PTPRC* expression for all spots, and observed an overall good correlation (**Fig. 3b**, Spearman cor.=0.7 and Pearson cor.=0.42), in contrast to the original expression (**Sup. Fig. 4a**). We also overlaid the differences between CD45 IF and *PTPRC* gene expression for each spot with respect to their physical locations. As is shown in **Sup. Fig. 4b**, Sprod has indeed significantly reduced the deviances between CD45 IF intensities and *PTPRC* gene expression for the majority of spots. We performed the same analyses with SAVER and scImpute as comparisons^6,7^, which are software tools for drop-out removal in regular scRNA-seq data. **Fig. 3c** showed that SAVER and scImpute only achieved modest improvement in terms of correlation. In addition, another control analysis with Sprod was performed by randomly permuting the image/spatial information. This “scrambled” control resulted in a very low correlation between CD45 IF intensities and the “denoised” *PTPRC* (**Fig. 3c**). This confirms that Sprod removed the noise through correctly borrowing information from external image/location information, rather than through merely smoothing of the expression data themselves.

**Fig. 3.** Validation of Sprod on real Visium and Slide-Seq datasets. (a) The Sprod-corrected *PTPRC* expression. Darker colors refer to higher expression. (b) Scatterplots showing the correlation between CD45 IF and *PTPRC* gene expression, corrected by Sprod. (c) The Spearman and Pearson correlations between CD45 IF and *PTPRC* expression, from the original expression data, Sprod-corrected expression data, expression data with drop-out removal performed by SAVER and scImpute, and the Sprod “scrambling” control. (d) The spatial IgD expression of the mantle zone marked in **Sup. Fig. 2** and **5**, for both the original Slide-Seq expression data and the Sprod-adjusted expression data. (e) Pearson correlations between IgD and CD3/CD20/CD1c, for each analysis group. (f,g) The Adjusted Exp.% of beads in the four bins of different sequencing qualities, in the Sprod-corrected expression data (f) and SAVER-corrected expression data (g), with the cutoffs to define the Adjusted Exp.% shown on the X axis. (h) Number of highly variable genes at different normalized dispersion cutoffs in uncorrected data, SAVER, Sprod “scrambling” and Sprod denoised data. Genes with average expression above 0.05 CPK and with normalized dispersion above the cutoffs were considered ‘highly variable’.

We next investigated a Visium human lymph node dataset. After the correction by Sprod, the adjusted IgD expression demonstrates a spatial pattern more concordant with the H&E stained images (IgD counts shown in **Sup. Fig. 5** and H&E stained image in **Sup. Fig. 2**). We highlighted one mantle zone in **Fig. 3d** (red circle in **Sup. Fig. 5** and the yellow circle in **Sup. Fig. 2**). The Sprod-corrected IgD expression formed a more distinctive ring-like pattern, compared with the original IgD expression (**Fig. 3d**), and is more consistent with the structures of mantle zones. To quantify the improvement of denoising by Sprod, we computed the expression correlations of IgD with several other genes. CD1c and CD20/MS4A1 are also markers of mantle zones and should be positively correlated with IgD^2^. In contrast, CD3 spans the perifollicular/interfollicular T cell regions and should be negatively correlated with IgD^3^. For the Sprod-corrected expression, CD1c/CD20 showed a much stronger positive correlation with IgD, and CD3 also showed a clearer negative correlation with IgD, compared with the original expression data (**Fig. 3e** and **Sup. Fig. 6**). Again, comparative analyses with SAVER and scImpute were performed, and also the “scrambling” control, which validated the superiority of denoising by Sprod.

Finally, we applied Sprod in the Slide-Seq mouse brain dataset used in **Fig. 1**. As in **Fig. 1f**, we calculated whether the over-inflation of highly expressed genes still persists after the Sprod-correction from the beads of lower sequencing quality. Unlike the original Slide-Seq data (**Fig. 1f**), the Sprod-corrected data have almost equal probabilities of observing highly expressed genes across the whole range of cutoffs for the four bins of beads (**Fig. 3f**). We performed the same analysis with SAVER (scImpute fails due to large data size), and observed worse performance, in terms of removing artificial differences between the four bins (**Fig. 3g**). We calculated the percentage of cells with zero counts (drop-outs) for each gene, and showed (**Sup. File 2**) that Sprod has significantly reduced drop-outs in the Slide-Seq data. We also calculated the numbers of highly variable genes (HVGs) in the raw and denoised expression data (**method section**). This assay can likely reveal whether the denoising process just over-smoothed the gene expression, in which case, the gene expression will become uniform and there will be less HVGs. Across different cutoffs to define HVGs, Sprod yielded up to 5 fold more HVGs than the raw data or the denoised expression data from Sprod-scrambled and SAVER (**Fig. 3h**). Our following analyses demonstrated that these HVGs detected from the Sprod-corrected data are indeed biologically meaningful.

### Detection of Spatially Variable Genes is More Accurate after Sprod Denoising

The detection of spatially variable genes (*i*.*e*. genes whose expression demonstrate certain spatial patterns) is one of the most prevalent analyses for ST data. The mouse hippocampal Slide-Seq data^1^ were examined once again. The hippocampus comprises several regions, including Cornu Ammonis (CA) 1, CA2, CA3 neurons, and dentate gyrus (DG). Stickels *et al* found several clusters of genes enriched in the dendritic region of CA1 by comparing beads in the proximal neuropil to the soma of neurons^1^. We tested whether the detection of spatially variable genes after the Sprod correction will be more meaningful. Importantly, we note that, due to the higher resolution of Slide-Seq and the nature of the neuronal cells in the mouse hippocampus regions, the Slide-Seq beads in this study captured different parts of the neurons, rather than different individual neurons. We normalized each bead’s expression data by per-bead library size, with the understanding that the normalized gene counts reflect the relative enrichment of mRNA transcripts within different regions of the neurons.

In **Fig. 4a**, the Slide-Seq beads that correspond to the soma, proximal neuropil, basal neuropil of the hippocampus neurons are highlighted in different colors, where these regions were identified using the same methods by Stickels *et al*. In **Fig. 4b**, the spatial expression of the *Camk2a* gene is displayed in the order of basil neuropil, soma, and proximal neuropil on the X-axis. **Fig. 4b** shows that there are severe drop-outs in *Camk2a*’s expression (dots concentrated at Y<0.1, 27.2% of all sequencing beads), and the beads unnaturally break into two groups by expression of *Camk2a*. The *Camk2a* expression in beads without drop-outs demonstrates a spatial gradient pattern of lower expression in the soma and higher expression in the neuropils. This spatial gradient reflects the fact that *Camk2a* is actively transported from the soma to the neuropils for local translation^12^. The other group of beads has *Camk2a* expression being strictly 0 and confounds the interpretation of the *Camk2a* expression pattern. To remove the drop-outs, we first used SAVER. However, as shown in **Fig. 4b**, SAVER only minimally recovered the non-zero expression of those beads with drop-outs (still concentrated close to Y<0.1, 27.1% of all beads), and the expression of *Camk2a* remains unnaturally dichotomized. In contrast, Sprod-corrected *Camk2a* expression has nearly completely eliminated the drop-out effects (3.5% of beads with expression<0.1) and the beads now demonstrate an overall consistent pattern of high expression of *Camk2a* in the neuropils and lower expression in the soma.

**Fig. 4.** Detection of spatially differentially expressed genes is more accurate after de-noising. (a) The Slide-Seq beads that were defined to be in the basal neuropils, soma, and proximal neuropils of the CA1 region, following the definition of Stickels *et al*. (b) The expression of *Camk2a* and *Hpca* in the CA1 region Slide-Seq beads, ordered along the soma-proximal axis. The X axis shows the location of the beads with respect to the soma. Results for the raw expression matrix, SAVER-adjusted matrix, and Sprod-adjusted matrix are shown. (c) Venn diagrams showing the overlap of the genes with differential expression detected from the raw expression data, the Sprod-adjusted data or the SAVER-adjusted data, with the genes that show dendritic enrichment from Tushev *et al* and Asinley *et al*. (d) Enrichment of GO pathways in the genes with stronger expression in the proximal neuropil regions, detected from the raw, the SAVER-corrected, and the Sprod-corrected expression data. The color of the circles refers to the P values of the GO analyses, and the size of the circles is proportional to the number of genes found in the pathway. (e) The percentages of dendritically enriched genes that are binding targets of each of the 7 RBPs or any of the 7 RBPs. (f) Sankey diagram showing the numbers of RBP binding targets in the differentially expressed genes identified from the raw, SAVER-corrected and Sprod-corrected data, for each RBP.

We also examined the expression of another gene, *Hpca*, whose expression is higher in the soma but lower in the neuropils. This gene’s expression also demonstrated a severe artificial dichotomization caused by drop-outs (**Fig. 4b**). SAVER is still unable to sufficiently remove the drop-outs. In contrast, in the Sprod-corrected data, drop-outs have been mostly removed. More importantly, the “raw” and “SAVER” results demonstrate a very sharp contrast between the soma bodies and the neuropils in terms of *Hpca* expression, with a sharp fall around X=100 (**Fig. 4b**). But for the “Sprod” expression data, there is a more natural and continuous gradient from the soma bodies to the neuropils. Finally, we also found that the genes whose expression showed positive or negative correlations with *Camk2a* and *Hpca* in the raw data now have enhanced positive or negative correlations in the Sprod-corrected data (**Sup. File 2**). These results prove that Sprod has indeed addressed drop-outs and corrected noises in the ST expression data in a biologically meaningful way, rather than merely numerically removing zero counts.

Next, we evaluated the effect of Sprod correction on the detection of spatially variable genes in a genome-wide manner. SpatialDE^13^, which is developed specifically for this purpose, was used to detect the genes with stronger expression in the proximal neuropil regions than the soma (**Sup. File 2**). In the uncorrected data, SpatialDE yielded 28 genes with differential expression at an FDR (False Discovery Rate)-adjusted P-value cutoff of 0.05, while in the Sprod-corrected data, SpatialDE identified 124 genes and, in the SAVER-corrected data, SpatialDE identified 222 genes. To validate whether these genes are in consistency with those previously reported, we cross-referenced Tushev *et al*^*14*^ and Ainsley *et al*^*15*^, who identified dendritically localized transcripts *via* microdissection and ribosomal-RNA enrichment, respectively. In **Fig. 4c**, we showed that the Sprod correction has greatly increased the sensitivity of differential expression analyses, while also retaining good specificity characteristics. The overlap between the differentially expressed gene set from Sprod-corrected data and Tushev *et al*+Ainsley *et al* achieved a Hypergeometric P-value of 6.98E-57 and a Log Odds Ratio(LogOR)=2.98). This is in comparison to the raw data (P-value=2.72E-19, LogOR=4.14), and SAVER-adjusted data (P-value=1.8E-5, LogOR=0.57), which indicates that SAVER correction has introduced too many false positives. This observation was confirmed by employing a simple Wilcoxon rank test to detect the differentially expressed genes (FDR-adjusted P-value <0.05, Fold-change >1.5). The Wilcoxon test yielded only 16 genes with differential expression in the uncorrected data, while in the Sprod-corrected data, the Wilcoxon test identified 42 genes, and surprisingly 2,752 genes for the SAVER-corrected data. The overlap between the differentially expressed genes and Tushev *et al*+Ainsley *et al* reached P-value=1.10E-26 and LogOR=3.82 for Sprod. For the raw uncorrected data, P-value=1.07E-11, LogOR=4.27, and for the SAVER-corrected data, P-value=2.99E-40, LogOR=0.73. Again, Sprod correction achieved the best balance of sensitivity and specificity for differential expression analyses.

We further evaluated the pathways in which these spatially variable genes are enriched. Examination of enriched GO pathways in the spatially variable genes from Sprod (compared with the raw data) shows that the corrected data lead to the discovery of more genes/pathways that are indicative of synapse functions (*e*.*g*. “neurotransmitter uptake”) or molecular transport in neurons (*e*.*g*. “axo-dendritic protein transport”) (**Fig. 4d**), consistent with the enrichment of these mRNA transcripts in the proximal neuropil regions. For the SAVER-corrected data, none of these pathways reached statistical significance. The dendritically enriched mRNAs are transported out of the soma bodies of the neurons, which is a process dependent on RNA binding proteins (RBPs)^16,17^. The CLIP-Seq technology is an experimental technique for profiling RNA-RBP interactions^18–20^. We downloaded CLIP-Seq datasets of several RBPs with known roles in dendritic mRNA transport or cytoplasmic stability from the ENCODE database^21^, which includes TDP-43^22^, SMNDC1^23^, IGF2BP1^24^, RBFOX^25^, FMR1^26^, CPEB^27^, and PUM2^28^. We calculated the percentage of the dendritically enriched genes that are binding targets of each RBP. In **Fig. 4e**, we show that the Sprod-corrected expression data yield dendritically enriched genes that are much more likely to be known targets of these RBPs according to CLIP-Seq, compared with the raw data and the SAVER-corrected data. We also performed an unbiased analysis by using all 122 unique RBPs’ CLIP-Seq data from ENCODE, and kept the top 25% most highly expressed RBPs according to this Slide-Seq dataset itself, plus the RBPs in **Fig. 4e. Fig. 4f** showed the number of RBP targets that can be identified within the significantly expressed gene sets from the raw, SAVER-corrected and Sprod-corrected data, for each RBP. The Sprod-corrected data yielded much higher numbers of these RBPs’ binding targets compared with the raw and SAVER-corrected data, even though the SAVER-corrected data possessed the largest number of differentially expressed genes.

Finally, in the proximal neuropil-enriched genes from the Sprod-corrected data, we discovered stronger over-representation of astrocyte terms (“astrocyte projection”, **Fig. 4d**), which conforms to the known existence of astrocytes and other glial cells around the neuropil regions of neurons^29–31^. Overall, our analyses prove that Sprod’s correction is critical for the accurate detection and interpretation of spatially variable genes and pathways.

### Sprod Facilitates Inference of More Accurate Spatial Pseudo-time and Cellular Communications

We further tested whether Sprod’s denoising improved the accuracy of the pseudotime inference. Pseudotime aims to infer the relative ordering of cells along a lineage based on the cells’ gene expression^32,33^. The breast cancer Visium dataset previously used in **Fig. 2e** was examined. We performed expression clustering of the spots and defined four tumor cell regions (**Fig. 5a**), with the remaining areas filled with stromal/immune cells. Given each Visium sequencing spot usually contains more than one cell, in order to ensure the inferred pseudotimes are more specific to the tumor cells, we removed the “contamination” from stromal/immune cells in the tumor region spots. To do this, we calculated a stromal/immune “contamination score” for each spot in the tumor region using a combined stroma/immune gene signature from Wang *et al*^*34*^ and Wang *et al*^*35*^. We removed spots with high “contamination scores”, and for the remaining spots, we also removed genes in this signature.

**Fig. 5.** Sprod facilitates inference of more informative spatial pseudo-time in ST data. (a) The four tumor regions (blue, green, orange, and red) that were extracted for pseudotime analyses, according to expressional clustering and concordance with the H&E stained slide. (b) The pseudo-times that were inferred by Slingshot for each of the four regions, respectively. Results for both the raw expression matrix (left) and the Sprod-corrected matrix (right) were shown. (c) An example gene, *CALML5*, with significant differential gene expression along the pseudotime trajectory, inferred by pseudotimeDE. Results for both the raw expression and the Sprod-corrected matrices, as well as for Slingshot-inferred pseudotimes and Monocle3-inferred pseudotimes were shown. (d) GSEA enrichment plots of the genes that demonstrate differential expression in the “early” and “late” (pseudotime) groups of spots in the left tumor region. (e) Boxplots counting the number of pathways that showed an absolute enrichment score >1.5 in the GSEA analyses in at least 3 out of all 4 tumor regions. (f) The pathways inferred from the Sprod-corrected data that showed an absolute enrichment score >1.5 in the GSEA analyses in at least 3 out of all 4 tumor regions, for both the Monocle-inferred and Slingshot-inferred pseudotimes. The enriched pathways from the raw expression data from the same analyses were excluded from this list.

Next, we predicted the pseudotimes of the tumor cells for each of the four regions, using SlingShot^32^ (**Fig. 5b**) and Monocle3^33^ (**Sup. Fig. 7**). To objectively and quantitatively prove that the pseudotimes predicted from the Sprod-corrected data are more biologically meaningful than those from the raw data, we examined the genes that demonstrate differential gene expression along the pseudotime trajectories (two example genes, *CALML5*^*36*^, and *CAPRIN1*^*37*^, shown in **Fig. 5c** and **Sup. Fig. 8**). Gene Set Enrichment Analysis (GSEA)^38^ was performed using the KEGG, GO, and cancer Hallmark gene sets (https://www.gsea-msigdb.org/gsea/msigdb/) (**Fig. 5d**) to detect the enriched pathways in the genes that demonstrate significant variation along pseudotime trajectories. We analyzed the concordance between the significant pathways detected by GSEA, among the four regions, as we reasoned that the pattern of tumor differentiation and phenotypic evolution should be similar within this small local region. As illustrated in **Fig. 5e** and **Sup. File 2**, the differentially expressed genes from each of the four regions are enriched in many more shared significant pathways for the Sprod-corrected data, than for the raw expression data. This holds true for both the Monocle3- and the Slingshot-inferred pseudotimes.

Next, the top common pathways between the four tumor regions from the Sprod-corrected data but not from the raw expression data were inspected more closely (**Fig. 5f**). Among those pathways, we identified the pathway of Hallmark_Epithelial_Mesenchymal_Transitionis, which is highly relevant for tumorigenesis^39^. More interestingly, many of these pathways are highly relevant for cell-to-cell communications, especially tumor-stroma/immune cell interactions, such as Collagen_Containing_Extracellular_Matrix and Integrin_Mediated_Signalling_Pathway.

Motivated by this and the fact that the pseudotimes of the spots appear to possess a difference between the centers of the tumor regions and the outer rims where the tumor cells interact with the stroma/immune cells (**Fig. 5b** and **Sup. Fig. 7**), CellChat was used to examine whether the interactions between the tumor cells and immune/stromal cells could be, at least in part, driving the differential transcriptomic status and thus impacting the pseudotimes of the tumor cells. We further divided (**Fig. 6a**) the tumor and non-tumor regions into 4 sections: region A (tumor cells not adjacent to stromal/immune (SI) cells), region B (tumor cells adjacent to SI cells), region C (SI cells adjacent to tumor cells), and region D (SI cells not adjacent to tumor cells). The classification procedure was described in the method section. Cellchat was deployed to infer cellular communications among the cells in these four regions. We hypothesize that long-distance tumor-SI cellular communications (B->D or C->A) should be substantially less active than the cellular communications at the boundaries of the tumor regions (B->C or C->B). Indeed, in the Sprod-corrected data, the number of significant ligand-receptor pairs inferred by CellChat is higher for the B->C communication than B->D, and higher for C->B than C->A (**Fig. 6b**). In contrast, the raw expression data yields a significantly reduced number of interacting pathways overall and more B->D than B->C interactions, which is less interpretable.

**Fig. 6.** Inference of cell-to-cell communication is more accurate with Sprod-corrected expression data. (a) The tumor and stroma/immune regions were both split into sub-regions that are either close or not close to the tumor-stroma/immune boundaries. These sub-regions were named as A, B, C, and D. (b) The numbers of CellChat-inferred significantly interacting pathways, in the raw and Sprod-corrected expression matrices. (c) The expression of *PD1* in the stroma/immune regions and *PD-L1* in the tumor regions, for the raw and Sprod-corrected expression matrices, respectively. (d) The close pairs of spots with one spot in the tumor region and another one in the tumor/stroma region. (e) The expression of *PD1* in the stroma/immune-side spots in the pairs of spots from (d), dichotomized by the expression of *PD-L1* in the corresponding tumor-side spots, and *vice versa*. Dichotomization was performed on the 75% percentile of *PD-L1* or *PD-1*. The bold lines in the boxes refer to median values.

In particular, we noticed that the pathway of *PD-L1* and *PD-1* in the significantly interacting pathways detected from the denoised expression data, but not in the original expression data. The role of the antagonizing interactions between *PD-L1* and *PD-1* in breast cancers has been well established^40–42^. In the denoised data, this pathway was observed to be significant in the direction of A->C and B->C, but with stronger confidence in B->C (CellChat Probability Score=2.21E-8, P value<1E-10) than in A->C (Probability Score=1.72E-8, P value<1E-10). We visualized the expression of the ligand *PD-L1*/*CD274* in the tumor cells and the receptor *PD1*/*PDCD1* in the stroma/immune cells in **Fig. 6c**, for both the raw and denoised data. Overall, the denoised data demonstrated a more obvious co-expression of *PD-L1* and *PD1* around the interfaces of the tumor-stroma/immune regions, compared with the raw expression. It is also evident that the expression of *PD-L1*/*PD-1* is not uniformly high along the interfaces, but rather possesses a local enrichment pattern. To objectively quantify the co-expression of *PD-L1*/*PD1* in the B/C regions, we defined close pairs of spots, with one spot in the B region and the other in the C region (**Fig. 6d**). In the pairs from the Sprod denoised data, the expression of tumor region *PD-L1* becomes much higher when neighboring stroma/immune regions demonstrate higher PD-1 expression (**Fig. 6e**, T test Pval.=5.6E-6). But when *PD-L1* becomes higher in the tumor cell regions, the expression of *PD-1* is only minimally higher in the stroma/immune regions (T test Pval.=6.1E-5, no difference in median values). This uni-directional observation is intriguing and also very reasonable, as we should anticipate tumor cells to up-regulate *PD-L1* expression in response to the cytotoxic pressure from PD1^+^ T cells, but not the other way around. In other words, this analysis revealed a causal relationship between the interaction of the PD-L1/PD-1 pathway. With the raw expression data, however, we cannot make this observation (**Fig. 6e**, Pval.=0.005 and 0.007, no difference in median value in either test). Overall, our above analyses indicated that Sprod also enabled more accurate inference of cell-to-cell communications.

## DISCUSSION

We developed Sprod to impute accurate gene expression in ST data. The existence of extensive noises were demonstrated in data from different ST technologies, which will impact downstream analysis severely and result in significant biases and misleading conclusions. Sprod took a novel approach of leveraging the physical location and matched imaging data of ST to remove such noises, rendering analysis and interpretation of ST data much more robust and accurate. Technically, the location/imaging similarity graph is obtained by an innovative sparse graph construction method based on a probabilistic density-based approach. This modeling strategy can tolerate high-dimensional data noise, preserve the pairwise metric, and integrate the imaging and positional features in a unified framework^43^. This allows Sprod to denoise (drop-outs and other noises) the expression of spots/beads in a dynamic neighborhood structure defined by imaging and location information. We systematically validated Sprod and its performance was demonstrated to be superior to algorithms designed solely for the removal of drop-outs in scRNA-seq data. Sprod is computationally efficient and user-friendly, and it is capable of readily handling data generated from a wide variety of ST technologies and being seamlessly integrated into bioinformatics pipelines for spatial transcriptomic analyses.

Drop-outs, as in regular scRNA-seq data, are one source of the inaccuracies in the ST data. Concerns have been raised in the field that drop-out correction methods for scRNA-seq data may introduce over-smoothing and thus erroneous signals in the scRNA-seq data^44,45^. Since regular scRNA-seq data only provide the gene expression of individual cells, these drop-out removal methods inevitably have to only rely on the cells with similar expressions to correct the drop-outs. This essentially bootstraps the expression data, which leads to the over-smoothing among cells. For ST data, Visium, Slide-seq, or similar technologies provide the spatial locations and often imaging features associated with the spots/beads. Sprod took the conceptually innovative approach of leveraging such external information for reliable imputation, which can avoid over-smoothing. Besides, the latent graph building process in Sprod enforces a neighborhood constraint (R) in denoising, further preventing the potential problem of over-smoothing.

Sprod is a universal application that can be used with all ST technologies. However, there are differences between ST technologies, so considerations should be given to how to best apply Sprod for each technology. This study focused on Slide-Seq and 10X Visium data, between which one distinction is the level of resolution. Each bead in Slide-seq likely profiles a 10μm by 10μm region while each spot in Visium profiles a 55μm by 55μm region. The sequenced cells and the multicellular structures of the tissues, on the other hand, are also of various size scales depending on the tissue and cell types. These factors together determine the degree of spatial dependency between spots/beads. The R parameter (physical scope to examine for spatial dependency) should be set larger for tissue types and ST technologies that impart a weaker spatial dependency, to introduce fewer constraints on spatial dependency in the graph building process. On the other hand, 3-dimensional sequencing technologies are also emerging, such as STARmap^46^. Currently, STARmap is only available for a small number of genes. However, whole transcriptome 3-dimensional sequencing will most certainly become available soon. With little modifications, the graph building model employed by the current Sprod model can be easily extended to consider spatial dependency in the 3-dimensional space (example in **Sup. File 2**).

Despite the great advancements of ST technologies, the field must pay close attention to the quality issues of ST data, which are more challenging than regular scRNA-seq. Though the current study focused on Visium and Slide-Seq, Sprod is also applicable to newer techniques such as HDST^3^ and Seq-scope^4^ (we provide our results on Seq-scope in **Sup. File 2**). It is important to note that HDST and Seq-scope (and, most likely, future ST techniques as well) achieve much higher resolution than Visium and Slide-seq. Noises in the ST data are likely to be more prevalent, necessitating even more cautious preprocessing before drawing any conclusions. Overall, we envision Sprod, developed specifically for imputation of accurate ST gene expression, to become a key first step to empower ST technologies for biomedical discoveries and innovations.

## MATERIALS AND METHODS

### Overview of Sprod

Sprod takes the positional and image features as input, calculates a latent graph from them, and finally uses the latent graph to smooth out noise in the original ST expression matrix. In our study, the ST expression matrices were transformed to the CPM (or CPK) scales and log-transformed. For future users, pre-processing of expression matrices (use of either counts/CPM/RPKM/TPM, normalization, batch correction, log transformation) is assumed to be the responsibility of the users and conducted before Sprod analysis, as necessary. The Sprod model will only use the expression matrices as is. The final output includes the denoised expression matrix, and the graph of spatial/location similarity of the spots/beads. The core Sprod model was implemented in the R language, and image manipulation and input/output interfaces were implemented in the Python language. The detailed model description was provided in **Sup. File 1**.

### Image processing and feature extraction

The Python ‘skimage’ package (version 0.18) was used for all image processing and feature extraction operations. Immunofluorescence (IF) and H&E images were normalized into 8-bit format first. For H&E images, additional steps including stain separation (skimage.colors.separate_stains) and adaptive histogram equalization (skimage.exposure.equalize_adapthist) were used to ensure the channels in the normalized image are minimally correlated.

Sprod relies on two types of image features based on the input data type. For datasets with matching IF or H&E images, two sets of features were extracted. For both sets of features, the image region where the features were extracted from can be the spot itself (‘spot’) or a box covering both the spot and its neighboring regions (‘block’). We calculated the 20th, 30th, 40th, 50th, 60th, 70th, and 80th percentiles of intensity values among all the pixels in each sequencing spot/block in each channel. These values were used as the intensity features for Sprod. We also calculated six sets of Haralick’s texture features^47^, including contrast, dissimilarity, homogeneity, ASM, energy, and correlation, for each sequencing spot/block in each channel. Specifically, the ‘skimage.feature.greycomatrix’ were used to extract the texture features with ‘offset = [1]’ and ‘angles = [0, π/4, π/2, 3π/4]’. We enable the option of choosing block-*vs*. spot-level and intensity *vs*. texture image features to the user. For the datasets used in our study, our choices were shown in **Sup. Table 1**.

For datasets without matching images, such as those generated by the Slide-Seq, pseudo-images were created. These features were generated in the following steps. Highly variable genes were selected using the ‘scanpy.preprocessing._highly_variable_gene’ method (https://github.com/theislab/scanpy/blob/f7279f6342f1e4a340bae2a8d345c1c43b2097bb/scanpy/preprocessing/_highly_variable_genes.py). Umap transformation was applied on the normalized dataset with only the highly variable genes, and beads in the transformed data were partitioned into clusters using the Dirichlet process^48,49^. The pseudo-image features were then assigned by the possibility of each bead belonging to each cluster.

### Scaling up in large datasets

Sprod is fast for datasets of thousands of spots/beads. However, for large datasets with tens of thousands of spots, special operations must be performed so that Sprod can run smoothly. In this work, we employed a splitting-stitching scheme to facilitate large dataset processing. Each Slide-Seq dataset was randomly (not dependent on spatial location) divided into *n* (10 by default) equal-sized subsets, and this process was repeated *b* (10 by default) times. Sprod denoising was performed on each of the *n* x *b* subsets and the denoised results were concatenated. Each spot was exactly denoised *b* times, and the concatenated denoised data from the *n* sampling batches were averaged so that the randomness resulting from the sub-sampling was averaged out.

### Generation of simulation data

In the simulation analyses (**Fig. 2**), three matrices were simulated as the input data: the “Expression matrix” (*E*), “Spots_metadata” (*C*), and “Image features” (*IF*). *E* had 100 genes and 5,000 spots and comprised 282 spots of cell type A (3 clusters), 232 spots of cell type B (2 clusters), and 4,486 spots of cell type C. *C* is the X/Y coordinates of the spots. A Dirichlet Process clustering of the expression matrix was performed on the expression data to detect the cell types, and the cell type labels and their assignment probabilities formed the *IF* matrix, in the same way as Sprod created the “pseudo-images”.

In particular, the expression matrix, *E*, was a sum of three parts: (1) Expressional variation of cell types, *E*_*1*_. It was generated from a multivariate normal distribution with three different means corresponding to the three cell types, and the covariance matrix was chosen such that three clusters of cells are visually discernible in the first two components of a Principal Component Analysis transformation. (2) Expressional variation of spatial dependency, *E*_*2*_. Here, we reasoned that spots of the same cell types that were nearby should have more similar expressions. We simulated *E*_*0*_ from a multivariate normal distribution with mean of 0 and the covariance matrix calculated based on the exponential of the negative of the Euclidean distances between all spots. The covariance among different cell types is set to zero so spatial dependency only happens for spots of the same cell types. *E*_*2*_ was scaled so that its variation was comparable to *E*_*1*_. (3) White noises in expression, *N. N* was a matrix of white noises generated from independent normal distributions with a mean of 0 and equal variance, which is controlled at several different levels (**Fig. 2c**). We admit that the white noise approach could be overly simplistic, which is a potential caveat of the simulation. But it’s hard to obtain the real distribution of the noises. The summed expression matrix is then *E=E*_*1*_+*E*_*2*_+*N*. Finally, the *E* matrix was transformed by an exponential function and scaled such that its distribution mimics the distribution of typical ST data.

The details of our simulation, especially all the numeric settings, can be found in our simulation script made available at: https://github.com/yunguan-wang/SPROD.

### Hyperparameter optimization

We employed grid search for determining an optimal set of tuning parameters for the simulated dataset. We evaluated combinations of these possible values: R (0.04-0.24, step size = 0.01), K (3-10, step size = 1), U (250, 500, and 1000), Lambda (0.1-1, step size = 0.1, plus 5 and 20), and L_E (0.3125, 0.625, 1.25, and 2.5). R is the radius to define the neighborhood of the spots. K is the dimension of the latent space. Lambda is a scaling parameter to control clustering information. U is the perplexity of the tSNE-like distance function of the input image data. L_E is a penalty parameter adjusting for the relative weight between the original expression matrix and the information from the spots in the neighborhood on the graph. In the real data applications, the parameters were selected by referring to the best parameter set according to the simulation dataset, which was determined by grid search, and are adjusted based on the diagnostic plots (**Fig. 2e**). The parameters used in all datasets involved in this work are listed in **Sup. Table 2**.

### Defining tumor and immune/stroma spots/regions that are close neighbors (Fig. 6)

In the breast cancer 10X Visium dataset, the spots were clustered based on gene expression, and manually examined and merged to the tumor and stroma/immune regions. Four tumor regions were defined (left, mid, mid-right, top-right) based on the cluster and spatial information. The tumor spots and their gene expression were cleansed for stroma/immune cell contamination. Specifically, a stromal gene module and an immune gene module were defined by the union of the gene signatures for stromal and immune cells defined by Wang *et al*^*34*^ and Wang *et al*^*35*^. Each spot was given a score for the stromal cells and a score for the immune cells, based on the average normalized expression levels of the genes in each module. Then, the tumor region spots with the top 5% of the module scores were filtered out. For the remaining tumor region spots, these immune- and stromal-related genes were also filtered out.

In **Fig. 6a**, we further split the tumor regions into two subsets: tumor regions not close to immune/stromal cells (A) or close to immune/stromal cells (B). For each spot in the tumor region, we will draw a circle based on a radius of twice the size of the spacing between two neighboring spots in the Visium spot lattice. We will count the number of stroma/immune region spots in this circle. This spot will be classified as “B” if its closest neighboring spots in the stroma/immune regions are more than a given cutoff. The cutoff is 15 for the left region, 25 for the mid region, 20 for the top-right region, and 15 for the mid-right region. This cutoff is slightly adjusted to ensure a smooth appearance of the classified “A” and “B” tumor regions. The stroma/immune region spots were classified similarly into “C” (close to tumor region) and “D” (not close to tumor region).

In **Fig. 6d**, we examined all pairwise connections between the type B spots and type C spots and calculated their distances. When the distance between a type B spot and a type C spot is smaller than three times the minimum of all distances, the two spots were counted as a close pair.

### Statistical analyses

All computations are conducted in the R or Python programming languages. Umap and tSNE were conducted by the umap v0.2.7.0 or Rtsne v0.15 R packages. Drop-out removals were conducted by SAVER v1.1.2 and scimpute v0.0.9. For all boxplots appearing in this study, box boundaries represent interquartile ranges, whiskers extend to the most extreme data point which is no more than 1.5 times the interquartile range, and the line in the middle of the box represents the median. All figures were made using the Python ‘matplotlib’ and ‘seaborn’ packages or the R ‘ggplot2’ package. In **Fig. 3h**, highly variable genes were defined using the ‘pp.highly_variable’ function in the ‘Scanpy’ Python package. Briefly, average expression and dispersions of each gene were calculated, then the dispersions were normalized by the average dispersions of genes with similar average expressions, and finally, highly variable genes were defined as genes with average expression and normalized dispersions above the three selected cutoffs. In **Fig. 4**, functional enrichment analysis using hypergeometric tests was performed using the ‘gseapy’ package in Python. In **Fig. 5**, the immune/stroma-contaminating spots/genes were cleansed as described above. For the remaining tumor region spots, pseudotimes were inferred using the Slingshot v1.8.0 or Monocle3 R packages. To be comparable, the roots, which were assumed to be the cells with the “earliest” developmental stages by Monocle3 were chosen to be also the spots with the earliest pseudotimes in Slingshot, in each of the tumor regions. It’s important to note that, at least in the case of linear lineage which is most common, the pseudotime by itself only determines which cells appear in similar developmental stages but the cells with higher pseudotimes could actually emerge earlier or later on in “real-time”. The differential expression genes along pseudotime trajectories were detected by the ‘pseudotimeDE v0.9.0’ package.

Pre-rank GSEA analyses were conducted by the ‘clusterprofiler v3.18.1’ package. For GSEA, the 9,000 top most variable genes were extracted from each tumor region and analyzed, to include as many genes as possible, but also to avoid including too many lowly variable genes that introduce ties in differential expression comparisons and thus difficulties in the clusterprofiler computation. The Visium tumor region spots were divided into “early” and “late” by the median of their pseudotimes. The “ranks” of the GSEA analysis for the genes were calculated as the log fold change of gene expression between the “early” and “late” groups. In **Fig. 6**, the cell-to-cell communications were inferred by the CellChat v1.1.3 package.

## Data availability

The Visium datasets are obtained from the public 10X resources/datasets website: https://www.10xgenomics.com/resources/datasets. The IDs of the datasets are: human-lymph-node-1-standard-1-1-0, Human-ovarian-cancer-whole-transcriptome-analysis-stains-dapi-anti-pan-ck-anti-cd-45-1-stand ard-1-2-0, human-ovarian-cancer-targeted-pan-cancer-panel-stains-dapi-anti-pan-ck-anti-cd-45-1-standard-1-2-0, and human-breast-cancer-block-a-section-1-1-standard-1-1-0. The ID of the regular 10X scRNA-seq dataset used in **Fig. 1c** is 10-k-peripheral-blood-mononuclear-cells-pbm-cs-from-a-healthy-donor-single-indexed-4.0.0. The Slide-Seq data are available from the publicly archived data by Stickels et *al*^*1*^. Specifically, we used the Puck_200115_08 data from https://singlecell.broadinstitute.org/single_cell/study/SCP815/highly-sensitive-spatial-transcriptomics-at-near-cellular-resolution-with-slide-seqv2.

## Code availability

The Sprod software is available at: https://github.com/yunguan-wang/SPROD. The DOI is 10.5281/zenodo.6047752.

## FIGURE AND TABLE LEGENDS

**Sup. Fig. 1** Correlation between *PTPRC* RNA expression (targeted) and CD45 protein expression (IF). Red arrows mean that the results are limited to spots of higher quality. Yellow arrows mean that the single gene of *PTPRC* is replaced by a signature of *PTPRC* by including correlated genes.

**Sup. Fig. 2** Overlaying the un-corrected IgD gene expression on the H&E stained image in the 10X Visium human lymph node dataset. The whole slide is shown, with the three examples of **Fig. 1d** picked from this slide. The yellow circle marks the mantle zone to be highlighted in **Fig. 3d**.

**Sup. Fig. 3** Gene expression clustering of the beads in the mouse brain Slide-Seq dataset reflects the multi-cellular structures of mouse brain hippocampus.

**Sup. Fig. 4** Correction of noise in the Ovarian Cancer Visium dataset. (a) Scatterplots showing the correlation between CD45 IF and the original *PTPRC* gene expression. (b) Deviances between CD45 IF intensities and the expression levels of *PTPRC* (left: original, right: denoised). CD45 IF intensities and *PTPRC* expression values were normalized and distributionally warped to the same scale so they can be directly compared. The differences between CD45 IF and *PTPRC* on each spot are denoted by color. Red refers to small differences and green refers to larger differences.

**Sup. Fig. 5** Spatial IgD expression of the raw Visium data (left) and the Sprod-adjusted data (right). The red circles mark the mantle zone to be highlighted in **Fig. 3d**.

**Sup. Fig. 6** Spearman correlations between IgD and CD3/CD20/CD1c for the human lymph node Visium dataset. Results are shown for the original expression data, SAVER/scImpute-corrected data, the Sprod-corrected data, and the Sprod-corrected data with image/location information scrambled.

**Sup. Fig. 7** The pseudo-times that were inferred by Monocle3 for each of the four regions, respectively. Results for both the raw expression matrix (left) and the Sprod-corrected matrix (right) were shown.

**Sup. Fig. 8** An example gene, *CAPRIN1*, with significant differential gene expression along the pseudotime trajectory, inferred by pseudotimeDE. Results for both the raw expression matrix and the Sprod-corrected matrix, as well as for Slingshot-inferred and Monocle3-inferred pseudotimes were shown.

**Sup. File 1** Mathematical details of the Sprod model.

**Sup. File 2** Additional analyses involving Sprod.

**Sup. Table 1** Tuning parameters used in Sprod for each dataset

**Sup. Table 2** Dendritically enriched genes in the mouse brain that are identified *via* different methods

## FUNDING

This study was supported by the National Institutes of Health (NIH) [CCSG 5P30CA142543/TW, GX, YX, 1R01CA258584/TW, U01AI156189/TW, YX, R01DE030656/GX, R01GM141519/GX, R01GM140012/GX, U01CA249245/GX, R35GM136375/YX, 2P50CA070907/TW, YX, GX], National Science Foundation [NSF DMS-2009689/LW], and Cancer Prevention Research Institute of Texas [CPRIT RP190208/TW, RP190107/GX].

## ACKNOWLEDGMENTS

We acknowledge the ENCODE Consortium and the ENCODE production laboratories that generated the eCLIP datasets used in our study. We acknowledge Dr. Jane Johnson for providing input on the interpretation of the mouse Slide-Seq data.

## AUTHOR CONTRIBUTIONS

Y.W. and B.S. implemented the software and contributed bioinformatics analyses. L.W. and T.W. designed the model. M.C. provided input on the interpretation of the pathology analyses. Y.X., S.W., and G.X. provided input on the analyses and the writing. T.W. supervised the whole study.

## COMPETING INTERESTS

The authors declare no competing interests

## Notes

### Competing Interest Statement

The authors have declared no competing interest.

