## Supplementary material for "Sprod for De-noising Spatial Transcriptomics Data Based on Position and Image Information": Table S1

| Dataset | Technique | Image features | R | K | Lambda |
|---|---|---|---|---|---|
| **Simulation** | Simulation | Pseudo-image | 0.1 | 10 | 0.4 |
| **Ovarian Cancer** | Visium | Spot level texture | 0.5 | 10 | 0.4 |
| **Lymph Node** | Visium | Spot level texture | 0.08 | 10 | 1 |
| **Mouse brain** | Slide-seq | Pseudo-image | 0.025 | 5 | 10 |
| **Breast Cancer** | Visium | Spot level intensity | 0.1 | 10 | 15 |

* The Ovarian Cancer and Lymph Node datasets participated in the validation/benchmark studies in **Fig. 3**, where the gold-standard criteria directly or in-directly involve correlation with the intensities of the image features. To alleviate any remote concern of leaking of information, we chose the texture features for building the latent graphs. For the breast cancer study, we chose the intensity features, which are generally better for latent graph building, according to our experiences.

** For the Visium datasets, the choices of R, K and Lambda are generally closer to the simulation study (first row). This is expected, as the simulation study is modelled after typical Visium datasets. The parameters chosen for the Slide-Seq study, determined by our diagnostic plots, are more different from the best parameter set from the simulation study, which is anticipated as well.

*** The R for the Slide-Seq dataset is much smaller than the Rs of the Visium datasets. This is due to the higher resolution of Slide-Seq, which provided much more refined information regarding the multi-cellular structures. Even with smaller R, there are still many beads in each radius defined by R in this dataset, which is used for borrowing information across beads.
