## Supplemental file 1 for "Sprod for De-noising Spatial Transcriptomics Data Based on Position and Image Information"

### Problem Description

More and more sequencing technologies are being created to provide the expression information and the spatial location information of the tissue slides around single cell resolution. However, these methods have a lot of noises in the gene expression measurements (drop-outs as in scRNA-seq and others). Luckily the spatial and image information can help with noise removal, and we developed the Sprod model for this purpose. In the Sprod model, a similarity graph of spots/beads is built based on image and positional features (Phase I). Afterwards, in Phase II, the gene expression is de-noised based on the similarity graph.

### Input Notations

There are three matrices we need as the input

1) The expression matrix $E = [e_{n,m}] \in R^{N \times M}$, where there are $N$ spots (using the terminology of 10X Visium as example) and $M$ genes. The algorithm will take the expression matrix into calculation as is, so any transformation and normalization, if needed, should be performed beforehand.
2) The spot coordinate matrix $C = [c_{n,j}] \in R^{N \times 2}$, where (X, Y) coordinates of these spots are specified.
3) The matrix of image features from the H&E or IF images as $IF = [i_{nl}] \in R^{N \times L}$. We use $i_n$ to denote row vectors of the $IF$ matrix. Here a Python-based functionality is provided to extract a default set of image features. Users can also feed additional customized image features extracted from the images. In the case of pseudo-images, this input is left empty, and the algorithm will calculate the pseudo-images from the input expression matrix.

### Phase I. Build a similarity graph of spots based on image and positional features

#### 1. Model Description

We propose to formulate the graph-building problem based on image/positional features as a manifold structure learning problem. Let $Y = [y_{n,k}] \in R^{N \times K}$ be the embeddings in some latent space of the learned graph, where $K$ is a tuning parameter for the dimension of the latent space. We took a probabilistic approach to model the embedding space. Assuming that the density function is $p(Y)$, and the density functions of $p(y^k)$ with respect to columns $y^k$ are independent:

$$p(Y) = \prod_{k=1}^{K} p(y^k).$$

We also assume a prior distribution for the latent space embeddings, which is assumed to follow the standard normal distribution:

$$\pi(Y) = \prod_{k=1}^{K} \pi(y^k) = \prod_{k=1}^{K} N(0, I),$$

where $I$ denotes the identity matrix.

Since $Y$ is a matrix of random variables, the pairwise Euclidean distance between $y_{n_1}$ and $y_{n_2}$ is defined as the expectation over $p(Y)$:

$$E_{p(Y)}\left[\left|\left|y_{n_1} - y_{n_2}\right|\right|^2\right] = \sum_{k=1}^{K} \int p(y^k)\left(y_{n_1,k} - y_{n_2,k}\right)^2 dy^k .$$

We define a tSNE-type distance measure to represent the similarity between spots in the original image feature space:

$$p_{n_1|n_2} = I(n_1 \neq n_2)\frac{exp\left(-\frac{d(i_{n_1}, i_{n_2})}{2\sigma_{n_2}^2}\right)}{\sum_{n \neq n_2} exp\left(-\frac{d(i_n, i_{n_2})}{2\sigma_{n_2}^2}\right)},$$

where $d(x, y) = \left|\left|x - y\right|\right|^2$ and $\sigma_n$ is the bandwidth of the Gaussian kernels. $\sigma_n$ is set in such a way that the perplexity of the conditional distribution is equal to a tuning parameter of $\mu$. $\sigma_n$ is found by using the bisection method to solve

$$\mu = 2^{-\sum_{n' \neq n} p_{n'|n} log_2\left(p_{n'|n}\right)},$$

between any two spots, $n$ and $n'$.

To preserve the information of the similarity between spots, provided through the original image feature data, we aim to achieve:

$$\frac{p_{n_1|n_2} + p_{n_2|n_1}}{2} = exp\left(-\frac{d(y_{n_1}, y_{n_2})}{\lambda}\right),$$

where $\lambda$ is a tunable parameter for controlling the clustering information. With Taylor expansion, we have

$$\frac{p_{n_1|n_2} + p_{n_2|n_1}}{2} = exp\left(-\frac{d(y_{n_1}, y_{n_2})}{\lambda}\right) = \sum_{s=0}^{\infty} \frac{\left(-\frac{d(y_{n_1}, y_{n_2})}{\lambda}\right)^s}{s!}.$$

In addition, we aim to control the embedding space such that as many as possible pairs of $(y_{n_1}, y_{n_2})$ satisfy $0 \leq \frac{d(y_{n_1}, y_{n_2})}{\lambda} \leq 1$. Thus, in such cases, the following condition satisfies:

$$\frac{p_{n_1|n_2} + p_{n_2|n_1}}{2} \leq 1 - \frac{d(y_{n_1}, y_{n_2})}{\lambda} + \frac{1}{2}\left(\frac{d(y_{n_1}, y_{n_2})}{\lambda}\right)^2,$$

which partially preserves the information and requirement of the preceding formula. Upon rearrangement, this formula turns into:

$$\frac{d(y_{n_1}, y_{n_2})}{\lambda} - \frac{1}{2}\left(\frac{d(y_{n_1}, y_{n_2})}{\lambda}\right)^2 \leq 1 - \frac{p_{n_1|n_2} + p_{n_2|n_1}}{2}.$$

In fact, we adopt a stronger condition for simplicity:

$$\frac{d(y_{n_1}, y_{n_2})}{\lambda} \leq 1 - \frac{p_{n_1|n_2} + p_{n_2|n_1}}{2},$$

which is convenient because $d(y_{n_1}, y_{n_2})$ satisfy $\frac{d(y_{n_1}, y_{n_2})}{\lambda} \leq 1 - \frac{p_{n_1|n_2} + p_{n_2|n_1}}{2} \leq 1$ naturally.

Furthermore, to take advantage of the spatial dependency information in the spatial sequencing data and to avoid over-smoothing, we require that only the spots that are close to each other to be considered in the graph. The physical closeness is defined as $n(n_1, n_2) = I(0 < (c_{n_1,1} - c_{n_2,1})^2 + (C_{n_1,2} - C_{n_2,2})^2 \leq R^2)$, where $R$ is a tunable parameter defining the radius of each spot's neighborhood.

Take all together, we estimate the density of embedded points *via* regularized Bayesian inference given by

$$min_{p(Y),\{\xi_{n_1,n_2}\}} \int p(Y) log\left(\frac{p(Y)}{\pi(Y)}\right) dY + S \sum_{n(n_1,n_2)=1} \xi_{n_1,n_2},$$

$s.t., (1) E_{p(Y)}\left[\left\|y_{n_1} - y_{n_2}\right\|^2\right] \leq \lambda\left(1 - \frac{p_{n_1|n_2}+p_{n_2|n_1}}{2}\right) + \xi_{n_1,n_2}, \xi_{n_1,n_2} \geq 0$, for $n(n_1, n_2) = 1$; and (2) $\int p(y^k)dy^k = 1$, for all $k$.

$S$ is a tunable parameter specifying the penalty strength of the slack variables. We note:

$$\int p(Y)log\left(\frac{p(Y)}{\pi(Y)}\right) dY = \sum_{k=1}^{K} \int p(y^k)log\left(\frac{p(y^k)}{\pi(y^k)}\right) dy^k,$$

according to the independent assumption of the prior distribution.

#### 2. Objective Function

The optimization problem is convex. By introducing multiplier variables $\{\alpha_{n_1,n_2} \geq 0\}, \{\beta_{n_1,n_2} \geq 0\}$ and $\{\omega_k\}$, we formulate the Langrage function:

$$L\left(p(Y), \{\xi_{n_1,n_2}\}, \{\alpha_{n_1,n_2}\}, \{\beta_{n_1,n_2}\}, \{\omega_k\}\right)$$

$$= \sum_{k=1}^{K} \int p(y^k)log\left(\frac{p(y^k)}{\pi(y^k)}\right) dy^k + S \sum_{n(n_1,n_2)=1} \xi_{n_1,n_2}$$

$$+ \sum_{n(n_1,n_2)=1} \alpha_{n_1,n_2}\left\{E_{p(Y)}\left[\left\|y_{n_1} - y_{n_2}\right\|^2\right] - \lambda\left(1 - \frac{p_{n_1|n_2} + p_{n_2|n_1}}{2}\right) - \xi_{n_1|n2}\right\}$$

$$- \sum_{n(n_1,n_2)=1} \beta_{n_1,n_2}\xi_{n_1,n_2} + \sum_{k=1}^{K} \omega_k(\int p(y^k)dy^k - 1).$$

According to the Lagrangian duality, we derive the KKT conditions as:

(1) $\partial_{\xi_{n_1,n_2}} L = S - \alpha_{n_1,n_2} - \beta_{n_1,n_2} = 0, \forall n(n_1, n_2) = 1$

(2) $\partial_{p(y^k)} L = 1 + log\left(p(y^k)\right) - log\left(\pi(y^k)\right) + \sum_{a(n_1,n_2)=1} a_{n_1,n_2}\left(y_{n_1,k} - y_{n2,k}\right)^2 + \omega_k = 0, k \in 1, \ldots, K$

(3) $\alpha_{n_1,n_2}\left\{E_{p(Y)}\left[\left\|y_{n_1} - y_{n_2}\right\|^2\right] - \lambda\left(1 - \frac{p_{n_1|n_2}+p_{n_2|n_1}}{2}\right) - \xi_{n_1,n_2}\right\} = 0, \forall n(n_1, n_2) = 1$

(4) $\beta_{n_1,n_2}\xi_{n_1,n_2} = 0, \forall n(n_1, n_2) = 1$

(5) $\int p(y^k)dy^k - 1 = 0, k \in 1, \ldots, K$

From (2), we see that

$$log\left(p(y^k)\right) = log\left(\pi(y^k)\right) - \sum_{n(n_1,n_2)=1} \alpha_{n_1,n_2}\left(y_{n_1,k} - y_{n_2,k}\right)^2 - 1 - \omega_k,$$

so we have

$$p(y^k) = \pi(y^k)\, exp\left(-\sum_{n(n_1,n_2)=1} \alpha_{n_1,n_2}\left(y_{n_1,k} - y_{n_2,k}\right)^2\right) / \, exp(1 + \omega_k).$$

As a result, $p(y^k)$ follows a multi-variate normal distribution with zero mean and precision matrix as $Q = I + 4diag(A\mathbf{1}) - 4A$, where $A = \left[I(n(n_1, n_2) = 1)\alpha_{n_1,n_2}\right]^{N\times N}$. In other words, we have

$$p(y^k) \propto exp\left(-\frac{1}{2}(y^k)^T Q y^k\right).$$

Accordingly, we have the dual problem by substituting the above equation to the Lagrange function $L$:

$$min \sum_{n(n_1,n_2)} \alpha_{n_1,n_2}\lambda(1 - \frac{p_{n_1,n_2} + p_{n_1,n_2}}{2}) + \sum_{k=1}^{K} \omega^k$$

To obtain the dual variable $\omega^k$, following equation (5), we have

$$\omega_k = log \int \pi(y^k)\, exp\left(-\sum_{n(n_1,n_2)=1} \alpha_{n_1,n_2}\left(y_{n_1,k} - y_{n_2,k}\right)^2\right) dy^k - 1.$$

Further, we have analytic form for the Gaussian integral as

$$\omega_k = log\,(det\,(Q)\,) - 1.$$

After dropping terms irrelevant to $A$, the dual problem is written as

$$min\, f(A) := \sum_{n(n_1,n_2)=1} \alpha_{n_1,n_2}\lambda(1 - \frac{p_{n_1|n_2} + p_{n_2|n_1}}{2}) - \frac{K}{2} log\,(det\,(Q)\,),$$

where $Q = I + 4diag(A\mathbf{1}) - 4A, s.t.\, 0 \le \alpha_{n_1,n_2} \le S, \forall n(n_1, n_2) = 1$

In practice, it is in fact possible to specify metrics other than $1 - \frac{p_{n_1|n_2}+p_{n_2|n_1}}{2}$ to measure the similarity between different spots, as long as the metrics are positive and more similar spots have smaller values.

##### 3. Optimization Algorithm

The above problem will be solved by the L-BFGS-B algorithm. The gradient is calculated as, for $\forall n(n_1, n_2) = 1$,

$$\partial_{\alpha_{n_1,n_2}} f(A) = \lambda \left(1 - \frac{p_{n_1|n_2} + p_{n_2|n_1}}{2}\right) - 2K(Q^{-1}_{n_1,n_1} + Q^{-1}_{n_2,n_2} - 2Q^{-1}_{n_1,n_2}).$$

Note that $\alpha_{n_1,n_2} = \alpha_{n_2,n_1}$ is treated as one single variable. When $\lambda$ is too large, $f(A)$ has a trivial solution of $A = 0$. Several trials may be taken to choose the largest possible $\lambda$ that does not lead to the trivial solution but can also impose enough strength of penalty.

#### Phase II. Spatial denoising based on the image/position graph

##### 1. Model Description

We obtain the optimal solution to $f(A)$ as $\hat{A}$, and we normalize the resulting graph such as:

$$g(n_1, n_2) = I(n(n_1, n_2) = 1)\frac{\alpha_{n_1,n_2}}{S}.$$

Recall that the original expression matrix is $E = [e_{n,m}] \in R^{N\times M}$, where there are $N$ spots and $M$ genes. The de-noised expression matrix is denoted as $D = [d_{m,n}] \in R^{N\times M}$, which is obtained through the following model. This model essentially computes a weighted average between the original expression matrix and the information borrowed from neighboring spots through the similarity graph:

$$min_D g(D) := ||D - E||_F^2 \ + \lambda_1 trace(D^T(diag(G\mathbf{1}) - G)D),$$

where $\lambda_1 > 0$ is a tunable regularization parameter. The first term is the mean square error to measure the difference between the original expression matrix and the de-noised expression matrix. By minimizing the error, we prevent the de-noised expression matrix deviating far away from the original expression matrix. The second term is the Laplacian smoothing term. It means that if two spots are neighbors on the graph $G$, their de-noised expression values of two spots should be similar. By the trade-off of the two terms, we can reach the expected de-noised expression matrix.

#### 2. Model Solution

The solution to the above model is easy to obtain. The gradient of the above loss function is:

$$g'(D) = 2(D - E) + 2\lambda_1(diag(G\mathbf{1}) - G)D,$$

and $g(D)$ is convex. So, the optimal solution is

$$\widehat{D} = [I + \lambda_1(diag(G\mathbf{1}) - G)]^{-1}E.$$
