## Supplemental file 2 for "Sprod for De-noising Spatial Transcriptomics Data Based on Position and Image Information"

### Sup. File 2: Additional Analyses Involving Sprod

#### Bead location and cell types of the simulation dataset

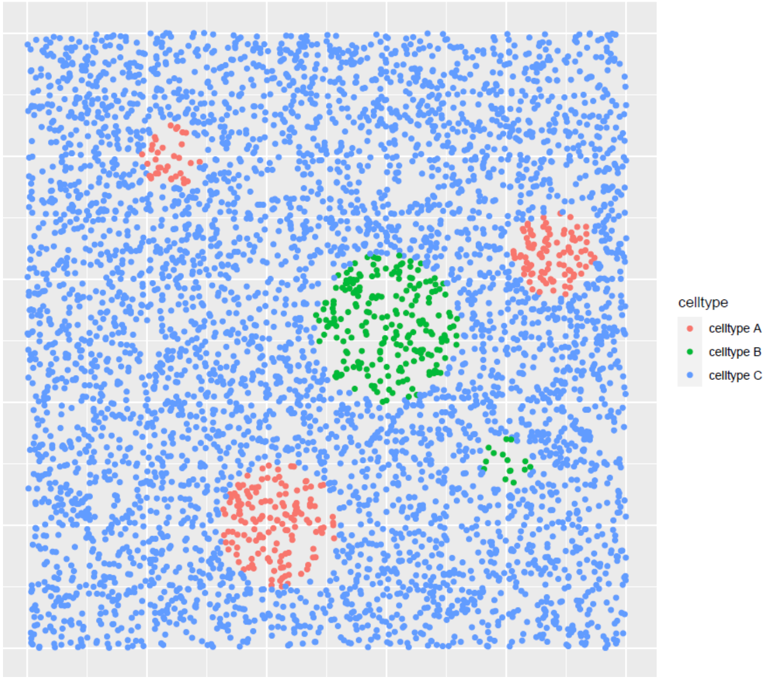


***SF2 Fig. 1*** *The locations of the beads and the cell types assigned to each bead in the simulation dataset*

#### Detection of spatially differentially regulated genes by SpatialDE

The SpatialDE algorithm detects genes that are spatially differentially regulated. It was run in both the raw and denoised Slide-Seq mouse hippocampus data using default parameters. Only cells in the soma, basal neuropil and proximal neuropil regions were analyzed. Since SpatialDE was agnostic of the predefined soma-to-proximal axis pattern, and we were only interested in genes enriched in the proximal neuropil region, additional analysis was performed to identify these genes. Briefly, genes were grouped into clusters that demonstrate similar patterns of regulation, and those clusters showing proximal neuropil region enrichment were picked *via* visual inspection. The figure below shows the genes (grouped into clusters) detected by SpatialDE to be spatially differentially regulated for the Sprod-corrected expression data. The original expression matrix was analyzed by SpatialDE in the same manner (data not shown). We extracted clusters 8, 10, and 13 in the results from the denoised expression matrix, and clusters 8 and 10 from the raw expression matrix.

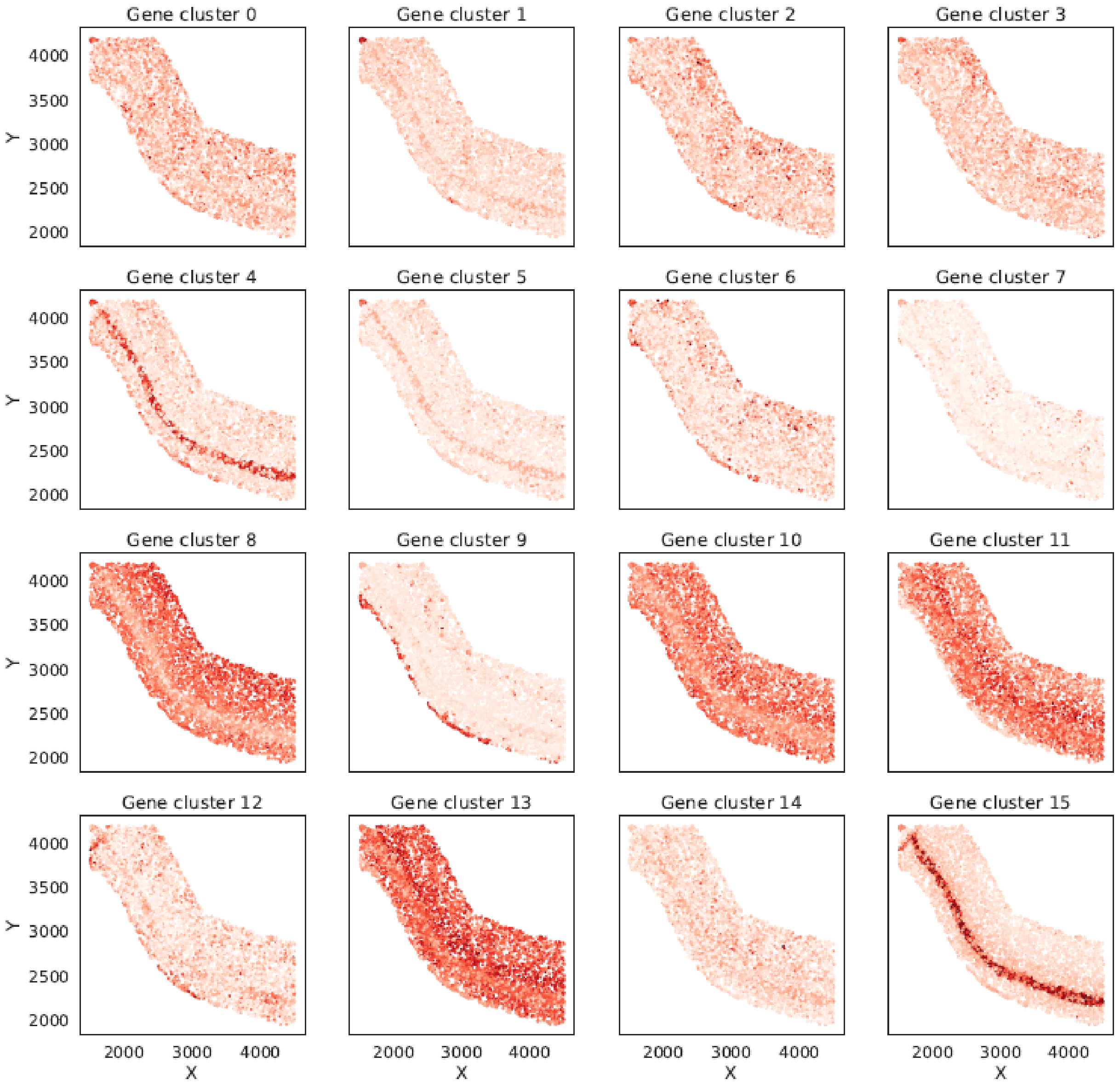


***SF2 Fig. 2*** *Genes (grouped into clusters) that are detected to be spatially differentially regulated by SpatialDE.*

**Sprod facilitates inference of more informative spatial pseudo-times in SEP data**

In **Fig. 5e**, we showed boxplots counting the number of pathways that showed an absolute enrichment score >1.5 in the GSEA analyses in at least 3 out of all 4 tumor regions. In this figure, we showed the results for “at least 2 out of all 4 tumor regions”, which is very similar to **Fig. 5e**.

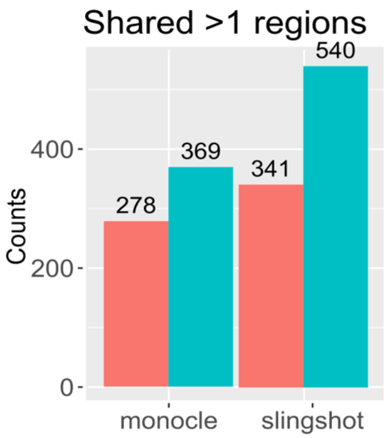


***SF2 Fig. 3*** *Boxplots counting the number of shared pathways that showed an absolute enrichment score >1.5 in the GSEA analyses in at least 2 out of all 4 tumor regions.*

**Statistical differences in pseudotimes between the boundaries and the centers of the tumor regions**

***SF2 Table 1*** *T test P values comparing the pseudotimes of the spots at the boundaries and in the centers of the tumor regions*

| | **Pval (monocle, raw)** | **Pval (monocle, denoise)** | **Pval (slingshot, raw)** | **Pval (slingshot, denoise)** |
|---|---|---|---|---|
| **Left** | <1E-8 | <1E-8 | <1E-8 | 3.9E-05 |
| **Mid** | 1.4E-06 | <1E-8 | 0.58 | <1E-8 |
| **Mid-right** | 0.024 | 0.00072 | 0.95 | 0.11 |
| **Top-right** | <1E-8 | <1E-8 | <1E-8 | <1E-8 |
